# Transcriptome-wide analysis of expressed resistance gene analogs (RGAs) in mango

**DOI:** 10.1101/2020.02.08.939736

**Authors:** Darlon V. Lantican, Cris Q. Cortaga, Anand Noel C. Manohar, Fe M. dela Cueva, Maria Luz J. Sison

## Abstract

Mango is an economically important fruit crop largely cultivated in the (sub)tropics and thus, is constantly challenged by a myriad of insect pests and diseases. Here, we identified and characterized the resistance gene analogs (RGAs) of mango from *de novo* assembly of transcriptomic sequences. A core RGA database of mango with 747 protein models was established and classified based on conserved domains and motifs: 53 nucleotide binding site proteins (NBS); 27 nucleotide binding site-leucine rich repeat proteins (NBS-LRR); 17 coiled-coil NBS-LRR (CNL); 2 toll/interleukin-1 receptor NBS-LRR (TNL); 29 coiled-coil NBS (CN); 4 toll/interleukin-1 receptor NBS (TN); 17 toll/interleukin-1 receptor with unknown domain (TX); 158 receptor-like proteins (RLP); 362 receptor-like kinases (RLK); 72 transmembrane coiled-coil domain protein (TM-CC), and 6 NBS-encoding proteins with other domains. The various molecular functions, biological processes, and cellular localizations of these RGAs were functionally well-annotated through gene ontology (GO) analysis, and their expression profiles across different mango varieties were also determined. Phylogenetic analysis broadly clustered the core RGAs into 6 major clades based on their domain classification, while TM-CC proteins formed subclades all across the tree. The phylogenetic results suggest highly divergent functions of the RGAs which also provide insights into the mango-pest co-evolutionary arms race. From the mango RGA transcripts, 134 unique EST-SSR loci were identified, and primers were designed targeting these potential markers. To date, this is the most comprehensive analysis of mango RGAs which offer a trove of markers for utilization in resistance breeding of mango.

## Introduction

Mango (*Mangifera indica* L.) is a globally popular fruit crop with economical and agricultural significance, especially in the (sub)tropics where it is largely cultivated. One of the constraints in mango production worldwide include a myriad of insect pests (e.g., oriental fruit fly, mango hopper, cecid fly, etc.) and diseases, (e.g., anthracnose, stem-end rot, scab, etc.) which can affect mango at different life stages thus significantly reducing fruit quality and yield (Bally, 2006). These are major barriers to the export market and can impede international trade as many pests and diseases in mango are important quarantine considerations.

Along the course of evolution, plants have generally developed mechanisms to recognize pathogen and insect attacks and to activate defense response against them (Dangl and Jones., 2001; Acevedo et al., 2015). The molecules derived from pathogens, insects, and plant cell damage (e.g., due to pathogen and insect attack) that trigger defense responses are generally referred to as microbe-/pathogen-/herbivore-/damage-associated molecular patterns (MAMPs/ PAMPs/ HAMPs/ DAMPs) (Acevedo et al., 2015; Choi and Klessig, 2016). These molecular patterns are being recognized by the pattern recognition receptors (PRRs) in the plant cell and trigger defense responses as these are perceived by the plant as “non-self” and indicative of pest attack (Pitzschke, 2013; Zipfel et al., 2006, Acevedo et al., 2015; Choi and Klessig, 2016), Hence, these molecules are generally considered as elicitors of plant defense responses and initiate the first line of defense, known as PAMP or DAMP-triggered immunity (P/DTI) (Jones and Dangl, 2006; Hu et al., 2018). The first layer of defense, however, is often overcome by plant pests through effector proteins, leading to effector-triggered susceptibility (ETS). To counteract this, plants can have a second line of defense known as effector-triggered immunity (ETI) (Boller and He, 2009; Pieterse et al., 2012). In ETI, due to host-pest co-evolutionary “arms race”, plants acquire a family of polymorphic and diverse resistance genes (*R* genes) which encode R proteins that recognize the attacker-specific effectors (Cui et al., 2015; Jones and Dangl, 2006) to correspond for a gene-for-gene interaction (Thompson and Burdon, 1992). This recognition results in intracellular signaling and switching ‘on’ of plant defense genes leading to plant resistance against pest attack.

Collectively, the PRRs and *R* genes are called resistance gene analogs (RGAs) which share conserved domains and motifs (Li et al., 2016). PRRs are predominantly categorized as receptor-like kinases (RLKs) and receptor like proteins (RLPs). RLKs have an extracellular sensing domain [either leucine-rich repeat (LRR) type or lysin motif (LysM) type], a transmembrane (TM) domain, and an intracellular kinase domain; while RLP possess similar domain structure except that it lacks an intracellular kinase domain (Tör et al., 2009). The R proteins are intracellular immune receptors (effector-recognition receptors) and majority of which belongs to nucleotide binding-site LRR (NBS-LRR) class (Gururani et al., 2012; Neupane et al., 2018). The R proteins contain certain domains or motifs such as serine/threonine kinases (STKs), LRRs, NBS, TMs, leucine-zipper (LZ), coiled-coil (CC) and toll/interleukin-1 receptor (TIR) (Li et al., 2016). Depending on the domain/motif architecture combinations, the subgroups of NBS-encoding proteins were designated as NBS, CC-NBS-LRR (CNL), TIR-NBS-LRR (TNL), CC-NBS (CN), TIR-NBS (TN), NBS-LRR (NL), TIR with unknown domain (TX), and other NBS-protein that may have chimeric domain/motif architecture. Since RGAs share conserved structural features, it is possible to predict them extensively to obtain a deeper insight on the underlying molecular defenses of the plant. RGAs have been widely studied and efficiently used for breeding of resistant crops against insect pests and diseases (Karthika et al., 2019). One way to predict RGAs in plants is through bioinformatics analyses of next-generation sequencing (NGS) genomic and transcriptomic data (Lantican et al., 2019; Neupane et al., 2018; Zhang et al., 2016; Karthika et al., 2019). With the advent of this technology, it is possible to unravel gene networks and develop molecular markers tagging economically important traits, reveal other molecular information such as intron-exon boundaries and presence of transposable elements, and discover novel biological processes (Goodwin et al., 2016).

The RGAs were first comprehensively studied on a genome-wide scale using the model plant *Arabidopsis thaliana* and since then, thousands of RGAs from numerous plant genomes have been identified (Meyers et al., 2003; Sekhwal et al., 2015). Despite the economical and agricultural importance of mango, especially in the tropics, its whole genome sequence (WGS) is yet to be available. Mango is believed to be an allotetraploid (4n=40) with genome size of around 450 Mb according to the on-going WGS effort (Chagné, 2015). Therefore, no detailed genome-wide analysis of mango RGAs has been performed, which is vital to facilitate breeding of resistant mango varieties. With the absence of such valuable resource, this study aimed to comprehensively identify and characterize the expressed RGAs of mango through *de novo* assembly of mango transcriptomes currently available in biological repositories. To our knowledge, this paper provides the most comprehensive identification and characterization of mango RGAs to date, which offer a trove of markers for direct utilization in resistance breeding of mango against insect pests and pathogens.

## Materials and Methods

### De novo transcriptome assembly

To obtain the expressed RGAs in mango, raw transcriptome reads were accessed and retrieved from public repository (NCBI; Table 1) and one directly from Hong et al. (2016). The raw read sequences were pre-processed by removing the adapter sequences and low-quality base score nucleotide sequences using Trimmomatic v0.36 (Bolger et al., 2014) with the following parameters: SLIDINGWINDOW: 5:30; LEADING:5; TRAILING:5; MINLEN:85. Each trimmed transcriptome FASTQ files was independently assembled *de novo* using Trinity pipeline (Grabherr et al., 2011) at default parameters. Upon assembly of the contigs, SuperTranscripts were then generated to provide an accurate representation of each expressed genes in the transcriptome assembly (Davidson et al., 2017). TransDecoder (Haas et al., 2013) was subsequently used to construct a polypeptide sequence database from the coding region of the SuperTranscripts based on nucleotide composition and open reading frame (ORF) length with default parameters.

**Table 1.** List of source transcriptome sequences and their predicted number of RGAs.

| Source SRA files | Mango variety | Tissue | NBS coding |  |  |  |  |  |  |  | RLP | RLK | TM-CC | Total |
| --- | --- | --- | --- | --- | --- | --- | --- | --- | --- | --- | --- | --- | --- | --- |
|  |  |  | NBS | CNL | TNL | CN | TN | NL | TX | Other |  |  |  |  |
| Hong et al., 2016 | 'ZILL' | Fr | 78 | 9 | 1 | 15 | 3 | 17 | 11 | 0 | 45 | 150 | 14 | 343 |
| SRR1448302 | 'KEITT' | Fr | 33 | 9 | 2 | 17 | 6 | 22 | 10 | 1 | 43 | 136 | 6 | 285 |
| SRR1561197 | 'KENT' | Fr | 20 | 0 | 0 | 3 | 1 | 1 | 4 | 0 | 15 | 68 | 12 | 124 |
| SRR1562177 | 'KENT' | Fr | 15 | 0 | 0 | 3 | 0 | 1 | 9 | 0 | 10 | 41 | 8 | 87 |
| SRR1956775 | 'AMRAPALI' | L | 40 | 1 | 0 | 9 | 1 | 2 | 11 | 0 | 10 | 21 | 6 | 101 |
| SRR2159471 | 'TOMMY ATKINS' | Fl, L, Fr, S | 74 | 7 | 1 | 15 | 3 | 18 | 12 | 2 | 78 | 223 | 21 | 454 |
| SRR2162836 | 'AMIN ABRAHIMPUR' | Fl, L, Fr, S | 71 | 10 | 1 | 11 | 4 | 23 | 18 | 1 | 76 | 216 | 22 | 453 |
| SRR2162878 | 'BURMA' | Fl, L, Fr, S | 24 | 0 | 0 | 3 | 0 | 2 | 11 | 0 | 37 | 112 | 11 | 200 |
| SRR2162889 | 'M. CASTURI "PURPLE"' | Fl, L, Fr, S | 85 | 6 | 1 | 7 | 2 | 14 | 19 | 1 | 67 | 177 | 20 | 399 |
| SRR2162908 | 'NEELUM' | Fl, L, Fr, S | 66 | 7 | 1 | 14 | 5 | 18 | 4 | 1 | 72 | 214 | 20 | 422 |
| SRR2162919 | 'THAI EVERBEARING' | Fl, L, Fr, S | 80 | 10 | 2 | 19 | 1 | 13 | 15 | 2 | 71 | 210 | 24 | 447 |
| SRR2162929 | 'TURPENTINE' | Fl, L, Fr, S | 67 | 6 | 0 | 10 | 5 | 15 | 14 | 2 | 71 | 186 | 23 | 399 |
| SRR2162953 | 'TURPENTINE', 'THAI EVERBEARING', 'NEELUM', 'M. CASTURI "PURPLE"', 'BURMA', 'AMIN ABRAHIMPUR' (Pooled) | Fl | 64 | 11 | 1 | 17 | 1 | 24 | 20 | 4 | 66 | 225 | 17 | 450 |
| SRR2165756 | 'TOMMY ATKINS' | S | 8 | 0 | 0 | 0 | 0 | 1 | 7 | 0 | 8 | 28 | 4 | 56 |
| SRR2736811 | 'ATAULFO' | Fr | 36 | 4 | 0 | 11 | 0 | 5 | 8 | 0 | 19 | 69 | 18 | 170 |
| SRR3192873 | 'AMRAPALI' | L | 89 | 7 | 1 | 18 | 4 | 15 | 15 | 4 | 49 | 193 | 16 | 411 |
| SRR3288569 | 'KEITT' | B | 75 | 4 | 1 | 17 | 2 | 13 | 17 | 0 | 61 | 157 | 21 | 368 |
| SRR3319054 | 'CHAUSA' | L | 68 | 7 | 1 | 19 | 6 | 15 | 12 | 2 | 60 | 187 | 17 | 394 |
| SRR3359450 | 'DASHEHARI' | Fr | 38 | 3 | 1 | 7 | 0 | 10 | 7 | 1 | 19 | 130 | 14 | 230 |
| SRR5966284 | 'AMRAPALI' | Fl | 32 | 6 | 0 | 16 | 5 | 11 | 14 | 1 | 45 | 212 | 21 | 363 |
| SRR5966306 | 'AMRAPALI' | Fl | 45 | 1 | 0 | 16 | 3 | 11 | 18 | 1 | 52 | 246 | 25 | 418 |
| SRR8449851 | 'CHOKANAN' | Fr | 53 | 1 | 0 | 5 | 2 | 8 | 10 | 0 | 20 | 88 | 12 | 199 |
| SRR8449858 | 'GOLDEN PHOENIX' and 'WATER LILY' (Pooled) | Fr | 103 | 10 | 2 | 19 | 6 | 23 | 15 | 2 | 56 | 168 | 12 | 416 |
| SRR8926025 | MANGO cv. 1243 | R; L; St, B,<br>Fl; Fr | 29 | 0 | 0 | 8 | 2 | 3 | 7 | 1 | 31 | 113 | 8 | 202 |
| **Total** |  |  | 1293 | 119 | 16 | 279 | 62 | 285 | 288 | 26 | 1081 | 3570 | 372 | **7,391** |

Fr: Fruit; Fl: Flower, L: Leaf, S: Seed; B: Buds; St: Stem; R: Roots

### RGA identification and classification

Mango transcriptome-wide resistance gene analogs (RGA) candidates belonging to Nucleotide Binding Site (NBS) and transmembrane-coiled-coil (TM-CC) containing proteins, and membrane associated receptor-like kinase (RLK) and receptor-like proteins (RLP) families were identified in the generated gene models from genome annotation using RGAugury (Li et al., 2016), an automated RGA prediction pipeline. The input protein sequences were initially filtered using BLASTp search against RGAdb database integrated in the pipeline using an *e*-value cut-off of 1e^−5^. The domain and motif of the initial set of candidate RGAs were detected using nCoils (Lupas et al., 1991), phobius (Kall et al., 2004), pfam_scan (Finn et al., 2010) and InterProScan 5 (Zdobnov and Apweiler, 2001) implemented within the RGAugury pipeline. A core set of RGAs were generated by concatenation and clustering (at 90% identity) of the identified RGAs from each polypeptide database using CD-HIT (Fu et al., 2012), and de-duplication based on BLAST description. The NBS-encoding proteins containing chimeric domain/motif architecture and designated by RGAugury as OTHER were manually checked for their domain/motif composition using Multiple Expectation Maximization for Motif Elicitation (MEME) (Bailey and Elkan, 1994) and MOTIF Search (https://www.genome.jp/tools/motif/).

### RGA characterization and annotation

Gene ontology (GO) annotations such as molecular function (MF), biological processes (BP), and cellular component (CC) were assigned to each protein represented in the constructed mango RGA core database using BLAST2GO package (Conesa et al., 2005). The homology of the predicted polypeptide sequences of each RGAs to existing entries in UniProtKB/SwissProt protein database were determined using BLASTp (with e-value of 1e^−5^). The mapped BLAST hits were then merged to InterProScan (Zdobnov and Apweiler, 2001) search output to produce the GO annotations.

### Profiling of mango RGA expression

RNA-Seq by Expectation Maximization (RSEM) (Li and Dewey, 2011) was used to provide insights on the expression profile of each RGA obtained from different tissues and diverse mango varieties. The pre-processed RNA-seq reads were independently mapped to the constructed core mango RGA reference transcriptome using Bowtie2 (Langmead and Salzberg, 2012). Transcripts per million (TPM) count values were then generated by RSEM from each paired-end reads. A comma-separated value (csv) file containing the RGAs and TPM values for each transcriptome data set was prepared and uploaded to Heatmapper (Babicki et al., 2016) for heat-map visualization of the RGA expression profiles.

### Evolutionary analysis

The FASTA amino acid sequences of the 747 core mango RGAs were used to construct a phylogenetic tree. Multiple Sequence Alignment (MSA) of the RGA sequences was performed using the CLUSTALW program (Thompson et al., 1994) with the following parameters: Gap Opening Penalty: 10; Gap Extension Penalty: 0.2. The phylogeny of these aligned sequences was reconstructed using Maximum Likelihood statistical method using IQ-TREE (Nguyen et al., 2015) with best-fit substitution model selected through ModelFinder (Kalyaanamoorthy et al., 2017). Based on the Bayesian Information Criterion (BIC) of the models, General ‘Variable Time’ (VT) matrix (Mueller and Vingron, 2004) with empirical amino acid frequencies (+F) and FreeRate (+R5) rate heterogeneity across sites (Yang 1995; Soubrier et al., 2012) was used to generate the tree. The resulting phylogenetic tree was validated with 1000 replicates of ultrafast bootstrapping (Hoang et al., 2017) and visualized using FigTree (v1.4.4) (Rambaut, 2018).

### Identification and design of EST-SSRs

The FASTA file corresponding to the RGA transcripts selected for analysis was uploaded into GMATA (Genome-wide Microsatellite Analyzing Toward Application) Software Package (Wang & Wang, 2016). SSR loci within the data set were identified using default parameters [Min-length (nt) = 2; Max-length (nt) = 6; Min. repeat-times = 5]. Consequently, oligonucleotide primer pairs were designed at regions flanking the identified SSRs using the Primer3 (Untergasser et al., 2012; Koressaar and Remm, 2007) integrated within the package. Default parameters were also used in primer design [Min. amplicon size (bp) = 120; Max. amplicon size (bp) = 400; Optimal annealing T_m_ (°C) = 60; Flanking sequence length = 400; Max. template length = 2000].

## Results and Discussion

### Identification of mango RGAs

Raw transcriptome sequences from diverse mango cultivars were retrieved, assembled, and the RGAs were identified using RGAugury, an integrative NGS bioinformatics pipeline that efficiently predicts RGAs in plants (Li et al., 2016). In general, this pipeline works by identifying first the domains and motifs related to RGAs such as NBS, LRR, serine/threonine and tyrosine kinase (STTK), CC, LysM, TM, and TIR. Based on the presence of combinations of these domains and motifs, the RGA candidates are then identified and categorized into four major families: NBS-encoding, TM-CC, and membrane associated RLP and RLK. The number of RGAs predicted from the sequence reads archive (SRA) files ranged from 56 to 454 (Table 1). In total, 7,391 RGAs were identified from the assembled transcriptome sequences (Table 1; Supplemental File 1). These RGAs were further clustered at 90% homology creating 2,892 clusters, followed by automated selection of a representative protein sequence per cluster. The representative protein sequences were described using BLAST and BLAST2GO, and proteins with duplicated BLAST description were removed retaining only unique/non-redundant protein type per RGA domain to generate 747 core RGAs with unique functions (Table 2, Supplemental Table 1 and Supplemental File 2). The core RGA proteins were classified as follows: 53 nucleotide binding site protein (NBS); 27 nucleotide binding site-leucine rich repeat proteins (NBS-LRR); 17 coiled-coil NBS-LRR (CNL); 2 toll/interleukin-1 receptor NBS-LRR (TNL); 29 coiled-coil NBS (CN); 4 toll/interleukin-1 receptor NBS (TN); 17 toll/interleukin-1 receptor with unknown domain (TX); 158 receptor-like proteins (RLP); 362 receptor-like kinases (RLK); 72 transmembrane coiled-coil domain protein (TM-CC); and 6 NBS-encoding proteins with other domains (Table 2).

**Table 2.**
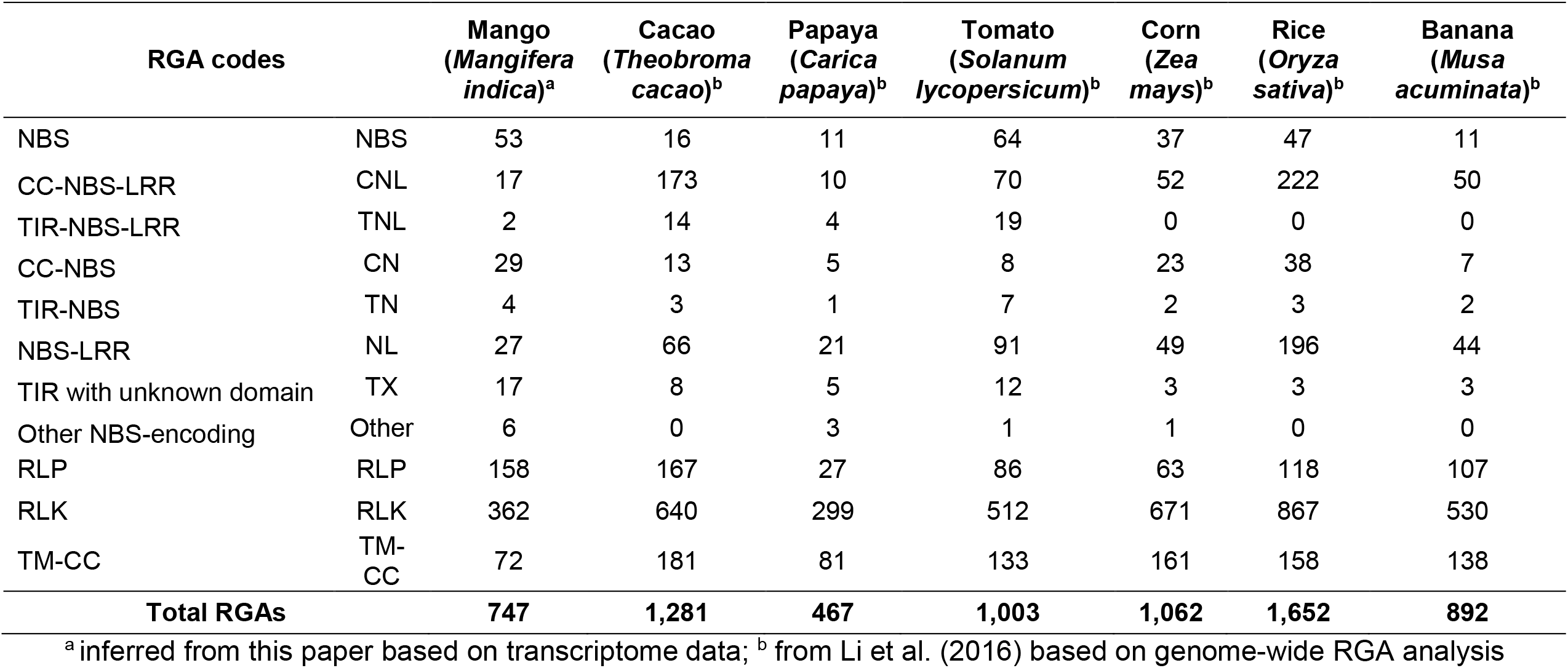
The distribution of core RGAs of mango and its comparison to other major crops.

Upon comparison to other important agricultural crops, as reported by Li et al. (2016), it appears that mango harbors more RGAs (747) than papaya (467) but has lower compared to cacao (1,281), tomato (1,003), corn (1,062), rice (1,652) and banana (892) (Table 2). Eventually, the RGA content and characteristics of a plant has been frequently associated with crop resistance (Sekhwal et al., 2015). Like most crops, RLKs constitute the largest group of RGAs in mango comprising almost 50% of the RGAs characterized, followed by RLPs, then the NBS-encoding proteins (Table 2, Fig. 1). In dicots such as mango, all NBS-encoding proteins are present (i.e., NBS, CNL, TNL, CN, TN, NL, TX, and other NBS-encoding proteins) (Table 2) but in monocots such as corn, rice, banana, the TNL protein is usually absent (Table 2) (Tarr and Alexander, 2009). It is hypothesized that after the divergence of dicots and monocots, the TNL genes might have been lost from the monocot lineage (Zhang et al., 2016). Six mango RGA models showed chimeric domain/motif architecture and were classified as ‘other’ NBS-encoding proteins (Table 2). These were manually checked using the MEME platform and appear to be NBS-encoding proteins containing TIR sequences and other uncharacterized domain (Supplemental Figure 1).

**Figure 1.**
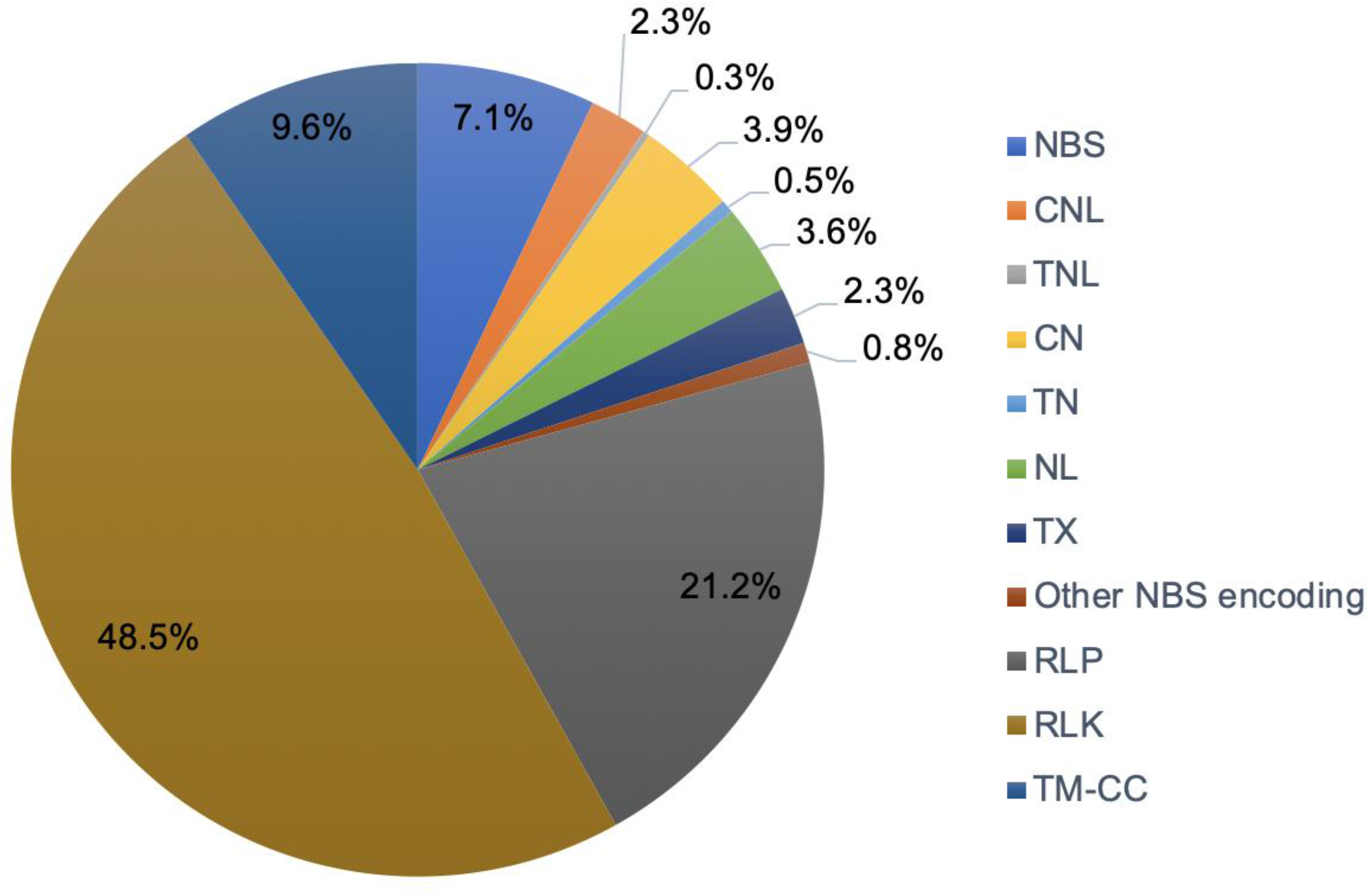
Distribution of the classifications based on conserved domains and motifs of core resistance gene analogs (RGA) detected in mango transcriptome *de novo* assemblies.

### Mango resistance (R)/defense proteins and expression profiles

BLAST analysis of the core mango RGA database revealed homology to a number of well-known resistance (R) /defense proteins against pathogens and insects (Table 3). Among these R proteins include RRS proteins (putative WRKY transcription factors), which confer resistance against *Colletotrichum higginsianum* and *Ralstonia solanacearum* (Narusaka et al., 2009; Saucet et al., 2015); RPP proteins (including At4g19530 and At4g19520 proteins) which confer resistance against downy mildew (*Peronospora parasitica*) (Parker et al., 1997; Sinapidou et al., 2004; Bittner-Eddy et al., 2000; Wan et al., 2019); the LRK10 proteins (all of which are of RLK domains) which provide resistance against leaf rust (*Puccinia triticina*) (Shiu and Bleecker, 2003; Feuillet et al., 1998); the RGA-blb and R1A-10 proteins which provide resistance against the devastating late blight disease (*Phytophtora infestans*) (Lokossou et al., 2010; Kuang et al., 2005); the RLM proteins which confer resistance against the fungal pathogen *Leptosphaeria maculans* (Staal et al., 2008; 2006); the Pik-2 and RGA5 proteins which confer resistance against blast disease (*Magnaporthe oryzae*) (Zhai et al., 2011; Okuyama et al., 2011); and many homologues of TMV resistance protein N which are responsible for tobacco mosaic virus (TMV) resistance (Dinesh-Kumar et al., 2000). The core mango RGA also contained numerous sequences with homology to various RPS proteins, including RPM1 and TAO1 proteins, which are important R proteins against the bacterium *Pseudomonas syringae*, a widely studied biotrophic pathogen (Eitas et al., 2008; Mackey et al., 2002; Warren et al., 1998; Kim et al., 2009).

**Table 3.**
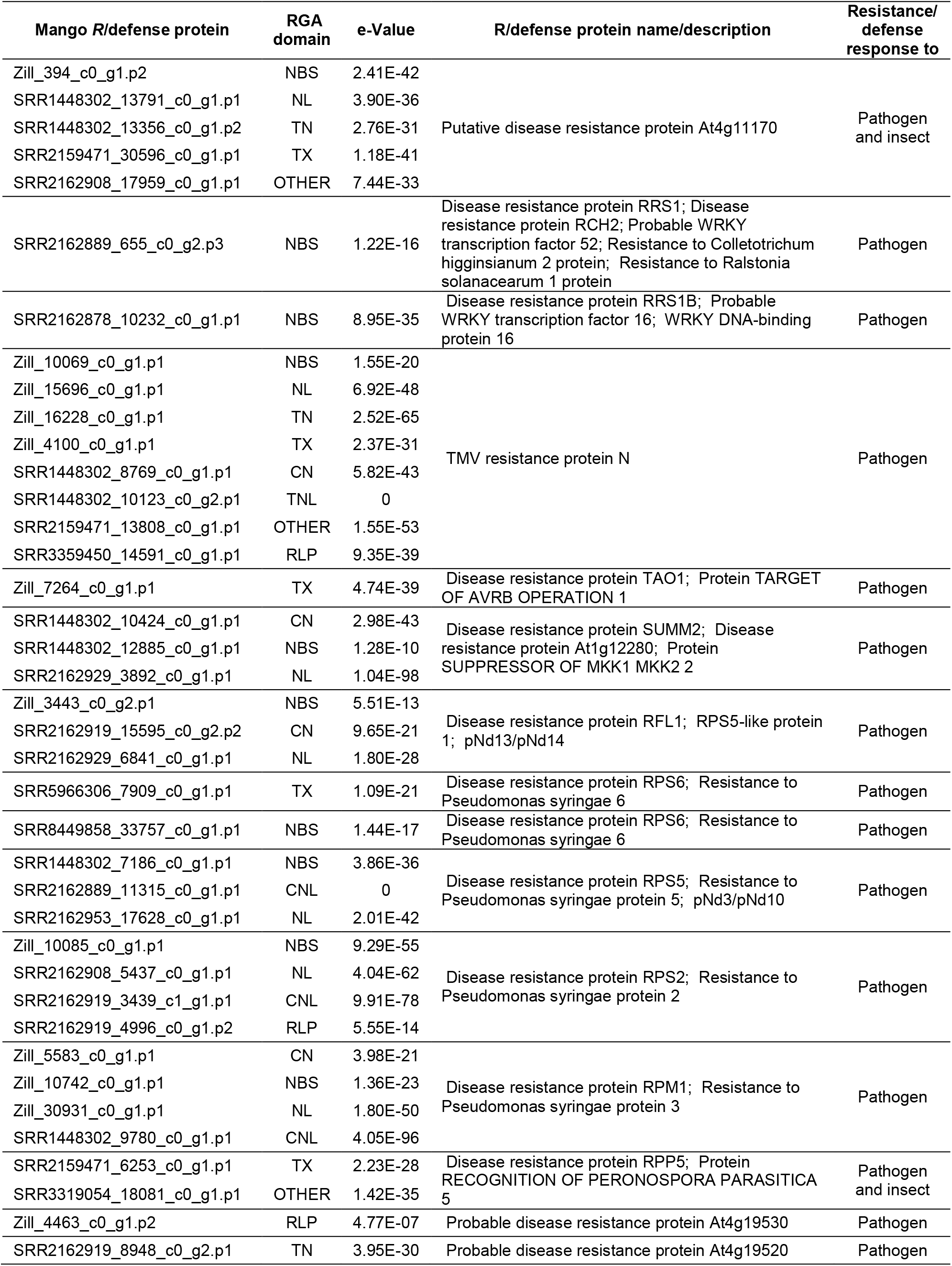
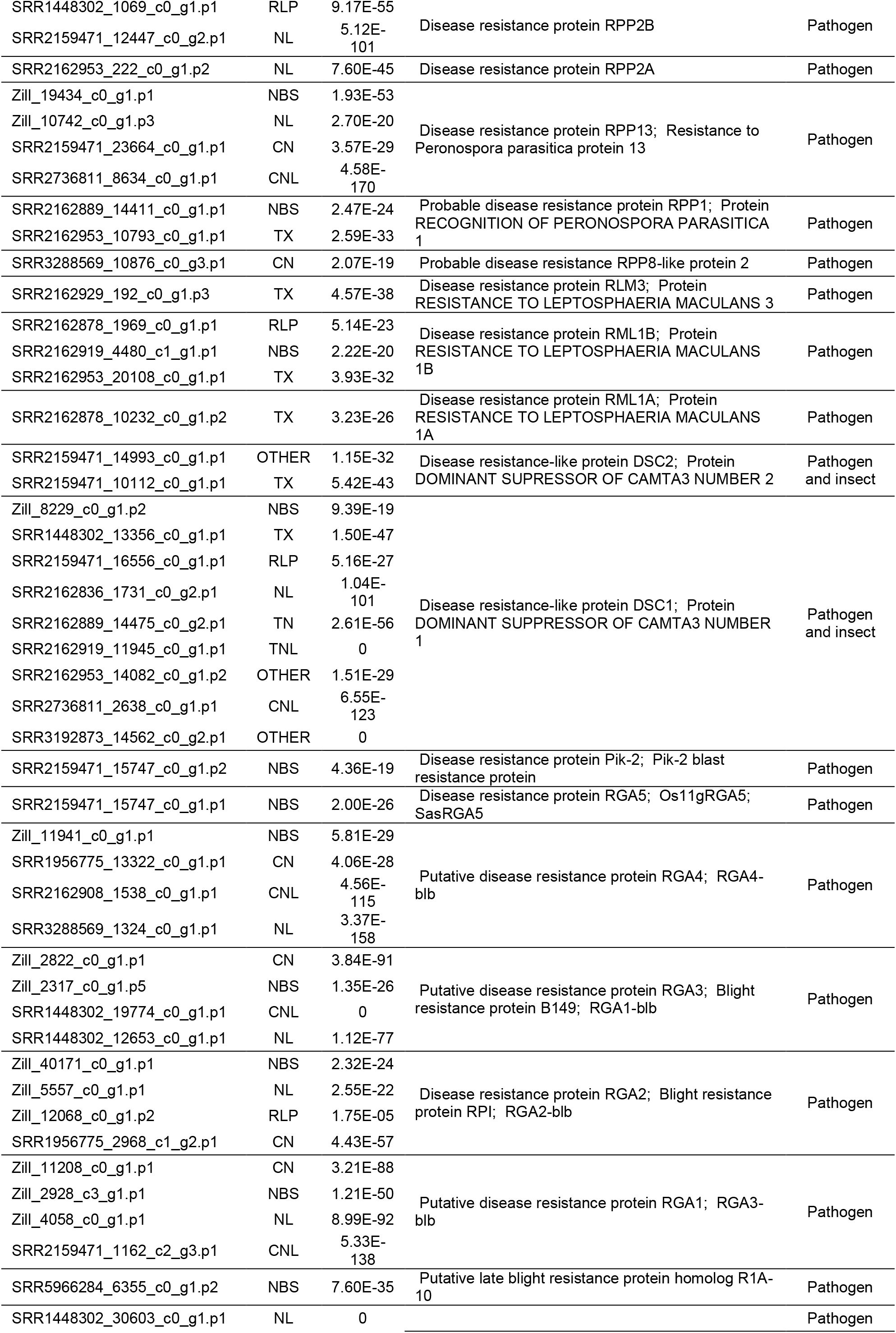
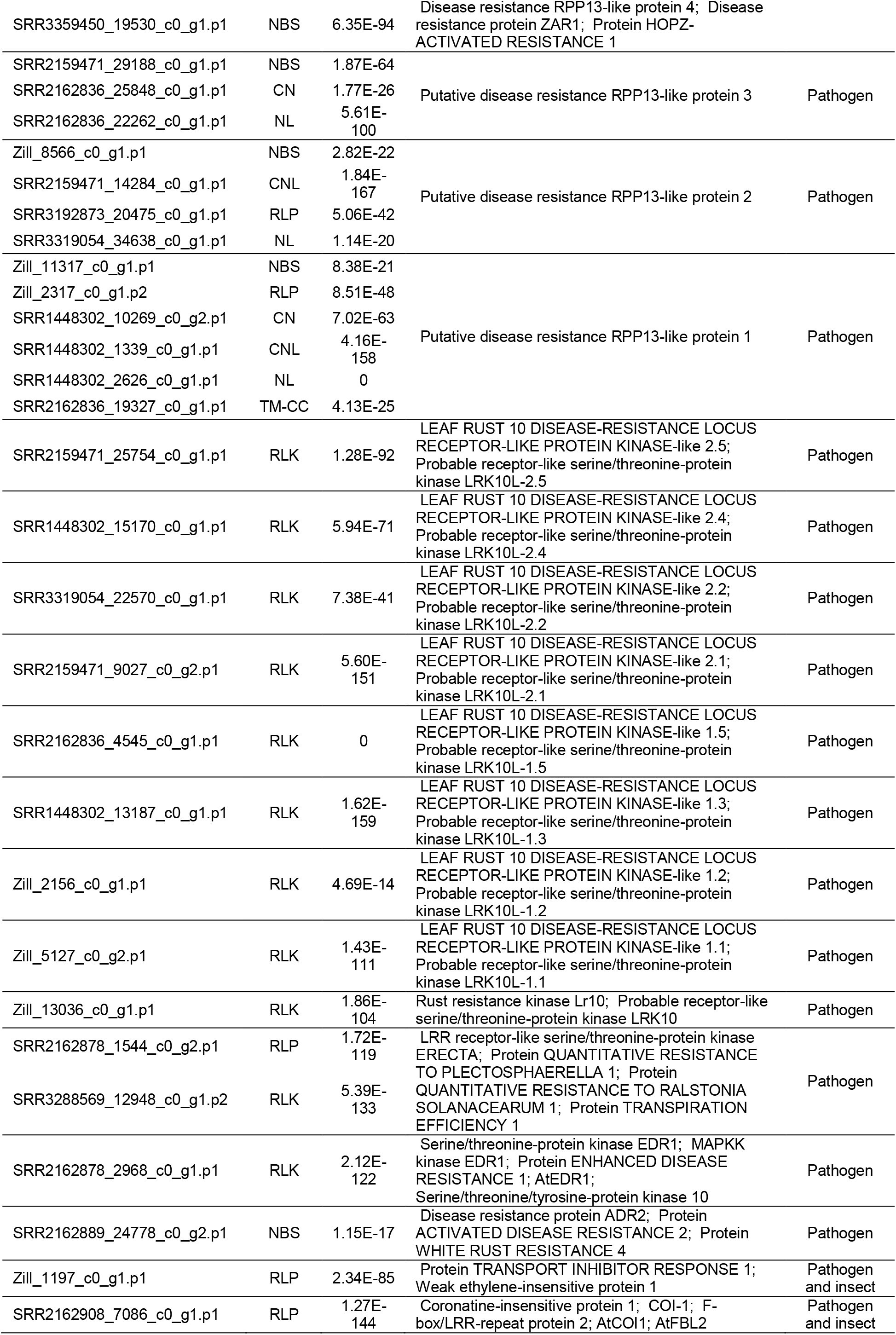
Some mango RGAs with homology to known Resistance (*R*)/defense proteins against pathogens and insects with homology to the entries in Uniprot/SwissProt database based on BLASTp hits.

Two RGA gene models showed homology to ERECTA protein which confer quantitative resistance against the necrotrophic fungus *Plectosphaerella*, and bacterial wilt caused by *Ralstonia solanacearum* (Sánchez-Rodríguez et al., 2009). This R protein is also responsible in the regulation of efficient transpiration in plant (Masle et al., 2005). Homology to the R protein EDR1 (ENHANCED DISEASE RESISTANCE 1), a MAPKKK serine/threonine-protein kinase, was also found which confer non-host resistance against *Colletotrichum gloeosporioides* in *Arabidopsis thaliana* through the induction of plant defensins expression (Hiruma et al., 2011). One mango RGA showed homology to the R protein ADR2 (ACTIVATED DISEASE RESISTANCE 2), an NBS-encoding protein that confers broad-spectrum resistance against many pathogens such as strains *of Pseudomonas syringae* and *Hyaloperonospora arabidopsis*, and full immunity to several races of the pathogen *Albugo candida*, the causal agent of white rust disease (Borhan et al., 2008; Aboul-Soud et al., 2009). This R protein is also reported to play a role in the response to UV stress (Piofczyk et al., 2015). Predicted mango RGAs homologous to the R-like proteins DSC 1 and 2 (DOMINANT SUPPRESOR OF CAMTA3) were also found which act as guard of CAMTA3, a negative regulator of immunity, during infection of pathogens (Lolle et al., 2017). Three NBS-encoding RGAs in mango showed homology to SUMM2 (SUPPRESSOR OF *mkk1 mkk2* 2) which acts as an R protein that becomes activated when the MEKK1-MKK1/MKK2-MPK4 cascade in the basal defense response is disrupted by the pathogen effector HopAI1 (Zhang et al., 2012; Kong et al., 2012).

Other putative resistance proteins abundantly found in mango showed homology to various DRL-coded (Uniprot/SwissProt ID) potential R proteins from *Arabidopsis thaliana* which have not been fully studied (Supplemental Table 1). Homologues to CHS and CHL proteins were also found in mango RGAs (Supplemental Table 1) which are important R proteins against chilling/low temperature and also confer resistance against bacterial infection such as *Pseudomonas syringae* (Zbierzak et al., 2013). Aside from resistance to pathogens, R/defense proteins against insects were also identified by filtering the gene ontology results with GO term “defense response to insect” (GO:0002213). The RGAs found include the At4g11170 putative resistance protein, RPP5, DSC1 and 2, TRANSPORT INHIBITOR RESPONSE 1 (TIR1), and CORONATINE INSENSITVE-1 (COI-1). TIR1 is an F-box protein and a major receptor of the plant hormone auxin and mediate in the auxin/indole acetic acid (IAA)-regulated signaling pathways involved in plant growth and development, as well as defense response against pathogens and insect pests (Dharmasiri et al., 2005). COI-1 is a jasmonic acid-isoleucine (JA-Ile) coreceptor and also an F-box protein involved in the JA signaling pathway which regulate plant defense against insects and pathogens, wound healing, and other vegetative and reproductive developmental functions (Xie et al., 1998). The plant hormone JA is conjugated to isoleucine (Ile) as a consequence of insect/pathogen attack, and JA-Ile binding with COI-1 stimulates degradation of JAZ coreceptor repressor proteins, thereby promoting the expression of JA-responsive genes (Pieterse et al., 2012). Further filtering of RGAs containing the GO term “response to insect” (GO:0009625) revealed 17 RGAs that putatively play vital roles in the defense pathways against insects (Table 4), all of which were characterized to possess RLK/RLP domains. The complete and detailed BLASTp analysis and description of the RGAs are provided in Supplemental Table 1.

**Table 4.** Other mango RGAs with homology to proteins in Uniprot/SwissProt database based on BLASTp hits and gene ontology analysis with “response to insects” annotation.

| Mango contig ID | RGA domain | e-Value | Description |
| --- | --- | --- | --- |
| SRR2162908_12681_c0_g1.p1 | RLK | 0 | Calmodulin-binding receptor kinase CaMRLK; |
| SRR8449851_23663_c0_g1.p1 | RLP | 6.40E-66 | Calmodulin-binding receptor-like kinase; AtCaMRLK; Protein MATERNAL EFFECT EMBRYO ARREST 62 |
| SRR5966284_6489_c0_g1.p1 | RLK | 4.04E-54 | Cysteine-rich receptor-like protein kinase 18; Cysteine-rich RLK18 |
| SRR2162953_2879_c0_g1.p2 | RLK | 7.43E-20 | Putative cysteine-rich receptor-like protein kinase 43; Cysteine-rich RLK43 |
| SRR5966306_10141_c0_g1.p1 | RLK | 3.68E-52 | Cold-responsive protein kinase 1 |
| SRR2162836_4413_c0_g3.p1 | RLK | 0 | Putative receptor protein kinase ZmPK1 |
| SRR2162836_3219_c0_g1.p1 | RLK | 0 | Probable LRR receptor-like serine/threonine-protein kinase RFK1; Receptor-like kinase in flowers 1 |
| SRR8449851_14359_c0_g1.p1 | RLK | 0 | Probable LRR receptor-like serine/threonine-protein kinase RKF3; Receptor-like kinase in flowers 3 |
| SRR2159471_39050_c1_g1.p1 | RLK | 0 | Probable receptor-like protein kinase At1g11050 |
| SRR3192873_14159_c0_g1.p1 | RLK | 0 | C-type lectin receptor-like tyrosine-protein kinase At1g52310 |
| SRR8449858_41086_c0_g1.p1 | RLK | 9.52E-66 | Probable LRR receptor-like serine/threonine-protein kinase At1g53420 |
| SRR1561197_1147_c0_g1.p1 | RLK | 8.21E-87 | Probable LRR receptor-like serine/threonine-protein kinase At1g56130 |
| SRR3192873_4970_c0_g1.p1 | RLK | 0 | Probable LRR receptor-like serine/threonine-protein kinase At1g07650 |
| SRR1448302_14204_c1_g1.p1 | RLK | 0 | Probable leucine-rich repeat receptor-like serine/threonine-protein kinase At3g14840 |
| Zill_8965_c0_g1.p1 | RLK | 1.47E-47 | Probable LRR receptor-like serine/threonine-protein kinase At1g53430 |
| SRR2159471_11990_c0_g1.p1 | RLK | 8.55E-62 | Probable LRR receptor-like serine/threonine-protein kinase At5g45780 |
| Zill_7246_c0_g2.p2 | RLK | 2.96E-117 | Probable LRR receptor-like serine/threonine-protein kinase At1g56140 |

Of all the mango varieties analyzed in this study, the three varieties that showed the lowest expression of RGAs were SRR8449851 (CHOKANAN), SRR3359450 (DASHEHARI), and SRR1562177 (KENT) (Fig. 2; Supplemental Table 2). On the other hand, the following showed the highest expressions of the RGAs characterized: SRR2162953 (pooled RNA from different varieties), SRR2162919 (THAI EVERBEARING), SRR2162878 (BURMA), SRR2162889 (M. CASTURI “PURPLE”), SRR2159471 (TOMMY ATKINS), SRR2162929 (TURPENTINE), SRR2162836 (AMIN ABRAHIMPUR), SRR2162908 (NEELUM) (Fig. 2; Supplemental Table 2). The relatively high expression of RGAs in these RNA-seq reads could be explained by the fact that they are pooled RNA samples from different varieties or from different plant tissues.

**Figure 2.**
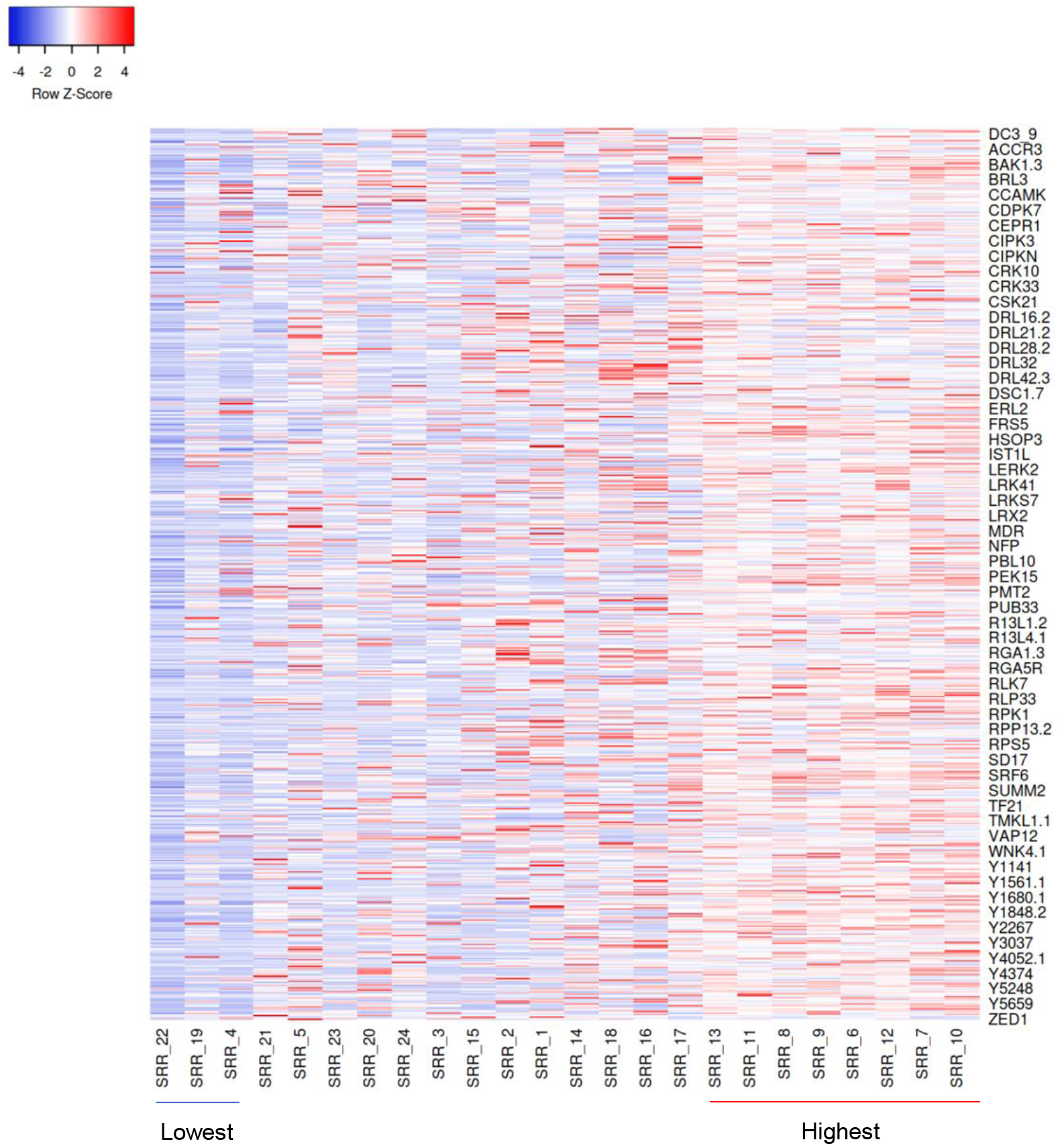
Resistance gene analogs (RGAs) expression profile from mango transcriptomes visualized as heatmap. Count values were normalized within samples as transcripts per million (TPM) using RSEM. Y-axis correspond to RGA gene IDs. X-axis correspond to RNA-seq raw read samples: *SRR_1. Hong et al., 2016 (ZILL), 2. SRR1448302 (KEITT), 3. SRR1561197 (KENT), 4 SRR1562177 (KENT), 5. SRR1956775 (AMRAPALI), 6. SRR2159471 (TOMMY ATKINS), 7. SRR2162836 (AMIN ABRAHIMPUR), 8. SRR2162878 (BURMA), 9. SRR2162889 (M. CASTURI “PURPLE”), 10. SRR2162908 (NEELUM), 11. SRR2162919 (THAI EVERBEARING), 12. SRR2162929 (TURPENTINE), 13. SRR2162953 (Pooled – TURPENTINE, THAI EVERBEARING, NEELUM, M. CASTURI “PURPLE”, BURMA, AMIN ABRAHIMPUR), 14. SRR2165756 (TOMMY ATKINS), 15. SRR2736811 (ATAULFO), 16. SRR3192873 (AMRAPALI), 17. SRR3288569 (KEITT), 18. SRR3319054 (CHAUSA), 19. SRR3359450 (DASHEHARI), 20. SRR5966284 (AMRAPALI), 21. SRR5966306 (AMRAPALI), 22. SRR8449851 (CHOKANAN) 23. SRR8449858 (GOLDEN PHOENIX and WATER LILY), 24. SRR8926025 (MANGO cv. 1243)*

### Gene ontology and functional annotation of mango RGAs

Gene ontology (GO) analysis and functional annotation of the core RGA database provided a broader knowledge on the major roles of these proteins and their cellular localizations. The molecular functions of the core RGAs are primarily associated with protein and nucleotide binding, and kinase activity (Fig. 3) as they are known extra- and intracellular binding receptors and relays defense signaling in the cells through a cascade of kinase activities (Rodriguez et al., 2010). The absence of an intracellular kinase in RLPs suggests that it relies/interact with other domains such as RLK-type receptors to relay the signalling from extracellular matrix to the intracellular component (Tör et al., 2009). Meanwhile, the biological processes of the RGAs are highly diverse, distributing the RGA sequences to a wide array of GO IDs/terms. The RGAs are mainly involved in protein (auto)phosphorylation during signal transductions in the cell, defense/immune responses to different biotic (insect pests and diseases) and abiotic (water deprivation, ozone, UV stress, etc.) stresses, and in various developmental processes (i.e., from embryonic to floral/pollen development) (Fig. 4). Numerous RGAs were also described to be involved in hormone-mediated signaling pathways and systemic acquired resistance which include the plant hormones brassinosteroids, abscisic acid (ABA), salicylic acid (SA), auxin, jasmonic acid (JA), ethylene, and gibberellic acid (GA). These plant hormones create a crosstalk to modulate defense signaling pathways and activate systemic resistance against a broad spectrum of pathogens and insect pests. As an example of this crosstalk, SA antagonizes JA to activate immune response against biotrophic pathogens (Williams, 2011). Likewise, JA antagonizes SA to activate two branches of immune response which also antagonizes each other: the ERF branch against necrotrophic pathogens (synergistic interaction with ethylene) and the MYC branch against insects (synergistic interaction with ABA) (Pieterse et al., 2012). In the cell, majority of the mango RGAs are localized in the plasma membrane, plasmodesma, cytosol, and nucleus which are primary sites for recognition of pathogen/insect invasion and effector proteins. In these cellular components, the RGAs functions to convert extracellular stimuli into intracellular responses to activate defense cascades and counteract the attack. In general, the results of GO analysis and functional annotation show that plant immune response is a consequence of the activity of a wide range of plant hormones, resistance/defense-related proteins, biochemical and developmental processes, etc. that aim to suppress pest attack thereby improving plant defense. The complete and detailed GO analysis and annotation of each RGAs are provided in Supplemental Table 1 and Supplemental Table 3.

**Figure 3.**
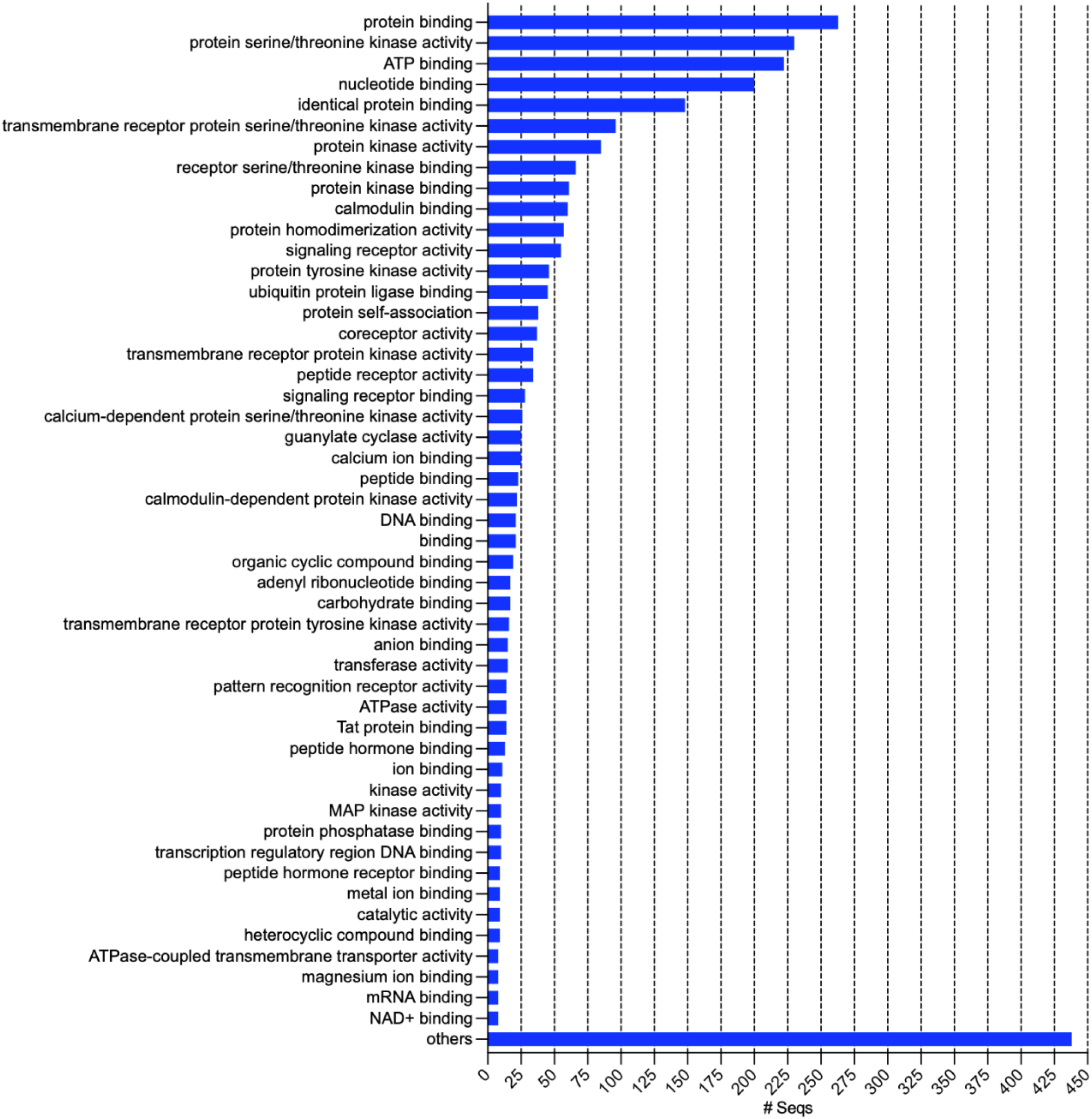
Gene ontology (GO) distribution of the expressed resistance gene analogs (RGAs) in mango based on molecular function (MF).

**Figure 4.**
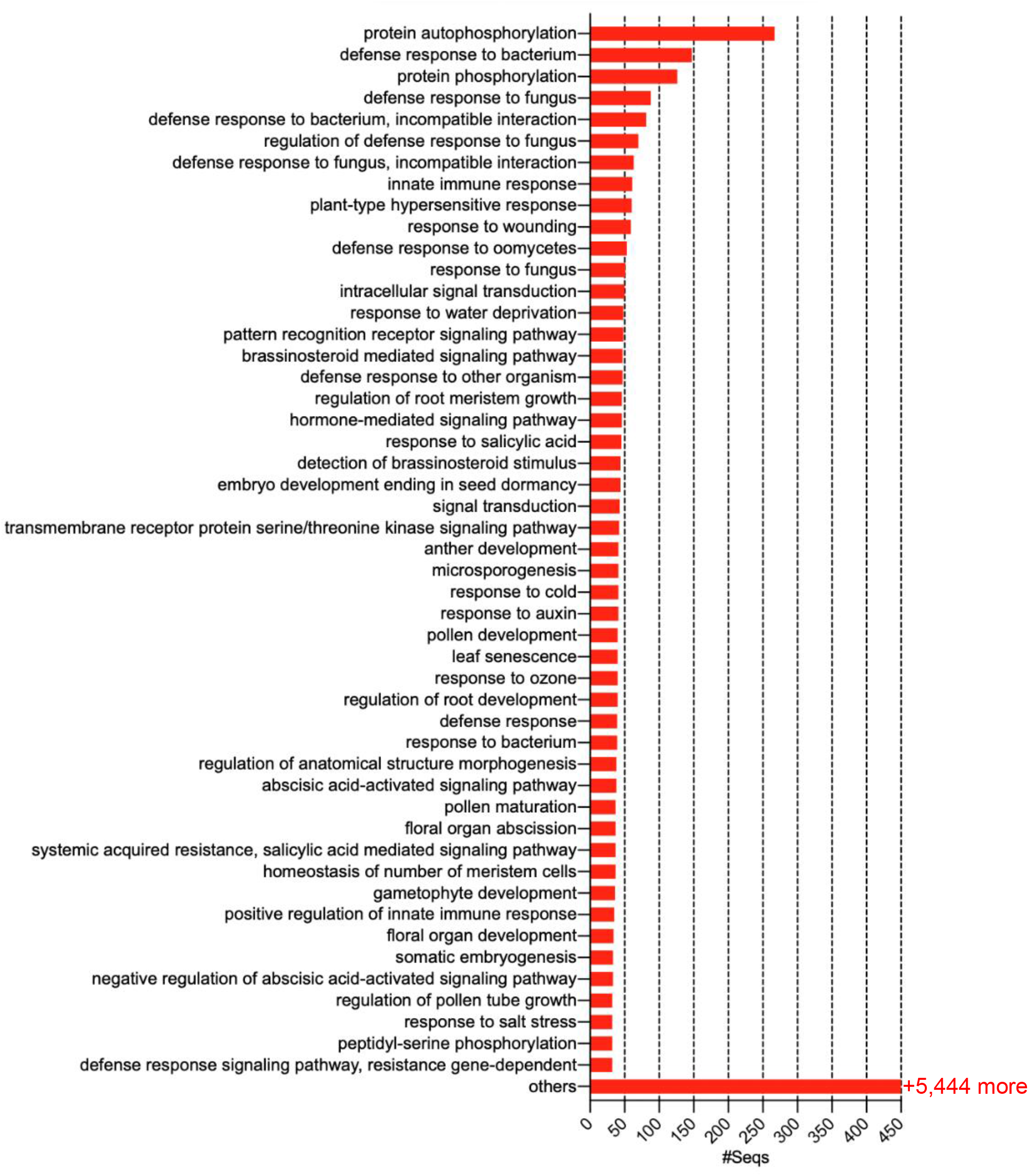
Gene ontology (GO) distribution of the expressed resistance gene analogs (RGAs) in mango based on biological processes (BP).

**Figure 5.**
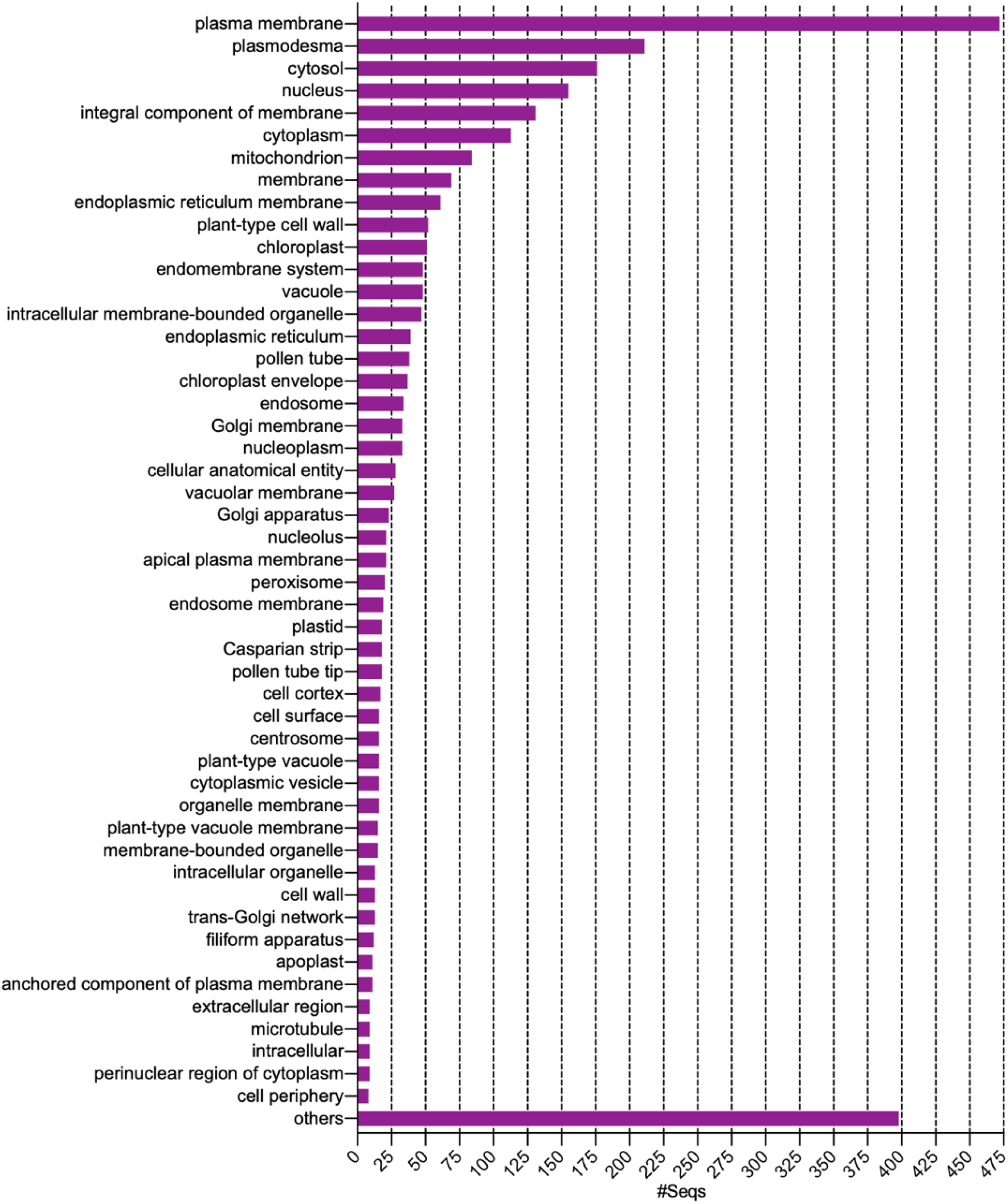
Gene ontology (GO) distribution of the expressed resistance gene analogs (RGAs) in mango based on cellular component (CC).

### Evolutionary relationships among mango RGAs

Using the evolutionary best-fit model selected according to Bayesian Information Criterion (BIC) (Supplemental File 3), a maximum likelihood phylogenetic tree was constructed using the 747 core mango RGA proteins to investigate their evolutionary relationships and diversity (Fig. 6; Supplemental File 4). It showed that clustering of RGAs based on domain classification is very apparent but mingled with subclades from other RGA domains. As shown in the tree, the mango RGAs have become highly diversified and are broadly clustered into 6 major clades with many subclades (Fig. 6). RGAs having TM-CC domain didn’t emerge as a separate cluster but formed subclades all across the tree suggesting that it shares similarity to all other RGA classes. The clade 1 is composed mainly of NBS-encoding proteins and other domains such as TM-CC and RLP. Notably, the clade 1 appeared to be the most distantly related RGA group among other clades. The clade 2 is predominantly composed of RLP proteins with subclades from all other RGA domains. This clade appears to be a sister clade to clade 3, which contains some RLP, TM-CC, and most especially, numerous RLK proteins which formed a major nested subclade indicating their close relationship with RLPs. The clades 4 and 5 appeared to be sister clades and are composed mainly of NBS-encoding proteins with subclades from all other RGA domains and most especially, from TM-CC domains. Rody et al. (2019) also observed that TM-CC sequences in sugarcane form widespread subclades across the phylogenetic tree and showed close relationship with NBS-encoding proteins. The clade 6 is composed predominantly of RLKs and shared close relationship to some other RGAs. In this clade, NBS-encoding proteins that shared close similarity related to RLKs were observed to form a major nested subclade. The NBS and RLKs are among the largest and diversified group of RGAs in plants (Hulbert et al., 2001; Meyers et al., 2003; Tör et al., 2009, Li et al., 2016), which also holds true in mango based on this study. The distribution of RGAs in the phylogenetic tree may indicate that their functions are highly divergent and form clusters that may be related to functions but not necessarily in protein sequence (Chang et al., 2002; Zhang et al., 2016). This could be the case in mango RGAs, especially the NBS-encoding proteins which are broadly distributed into 3 major clades (clades 1, 4 and 5) and 1 major subclade (nested in clade 6), as well as in RLKs which formed a major clade (clade 6) and a major subclade (nested in clade 3).

**Figure 6.**
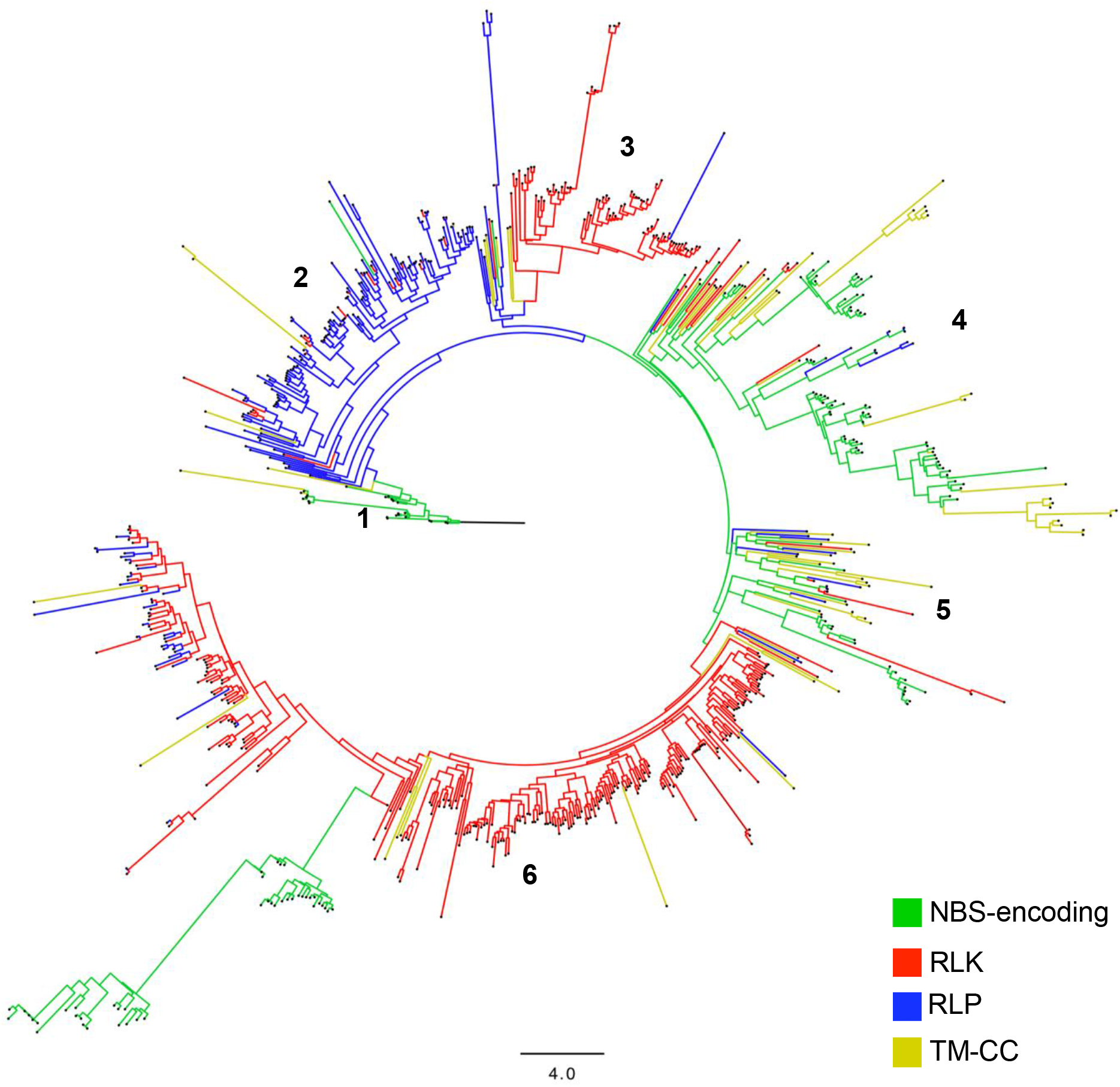
Maximum likelihood phylogenetic tree constructed using IQ-TREE from the sequence alignment of the core resistance gene analogs (RGAs) generated from *de novo* transcriptome assemblies of mango. Best-fit model was selected according to Bayesian Information Criterion (BIC) using ModelFinder (Kalyaanamoorthy et al., 2017). General ‘Variable Time’ (VT) matrix (Mueller and Vingron, 2004) with empirical amino acid frequencies (+F) and FreeRate (+R5) rate heterogeneity across sites (Yang 1995; Soubrier et al. 2012) was used to generate the tree, tested with 1000 replicates of ultrafast bootstrapping (Hoang et al. 2017). The numbers correspond to the clades.

Many factors have been attributed to the evolutionary pattern and diversity of RGAs in plants. One of which is the well-known co-evolutionary “arms race” model between the host plant and associated pests and diseases which drives the selective forces to overcome each other (Anderson et al., 2010; Edger et al., 2015; Boller and He, 2009). Other selection pressures such as climatic conditions (e.g. rainfall, temperatures, humidity, etc.) that favors pest and disease development have been correlated also to the diversity of RGAs (Sela et al., 2009). Whole-genome duplications (WGDs) and genomic reorganizations that occurred during ancient times, especially in angiosperms, have been associated to the expansion of RGA families and in creating new gene functions (Perazzolli et al., 2014, Chagné, 2015; Michelmore and Meyers, 1998). It is likely that in the evolutionary history of mango, it had undergone at least one round of polyploidization i.e., allopolyploidy which is a type of WGD via hybridization followed by genome doubling (Glover et al., 2016; Soltis and Soltis, 2009). These are some factors that could have influenced the complex phylogenetic structure of RGAs of mango, given also that the plant’s tropical growing condition provides a thriving environment to a plethora of insect pests and diseases.

### RGA-derived EST-SSRs of mango

Using the RGA transcripts (Supplemental File 5), a total of 151 EST-SSR (Expressed Sequence Tags – Simple Sequence Repeat) loci were identified. Among these, 134 loci yielded unique potential markers. Mostly di-repeat motifs (2-mer) were observed, accounting for 64.24% of the markers, followed by tri-repeats (3-mer) at 30.46%. The remaining 5.3% is composed of tetra-, hexa-, and penta-repeat motifs. AT repeat is the most represented motif among the di-repeats at 12.5%, followed by TC and TA at 11.26% each (Fig. 7). Meanwhile, AG/CT is the most abundant paired repeat motif among the SSR loci analyzed at 16.56% (Fig. 8). Data on the distribution of *R* gene motif-linked SSRs (Fig. 9) revealed that RLK has the most number of EST-SSRs designed with 67 markers, followed by RLP and TM-CC motifs at 34 and 18 markers, respectively. This distribution corresponds directly to the amount of *R* genes per category identified in the whole mango RGA transcriptome.

**Figure 7.**
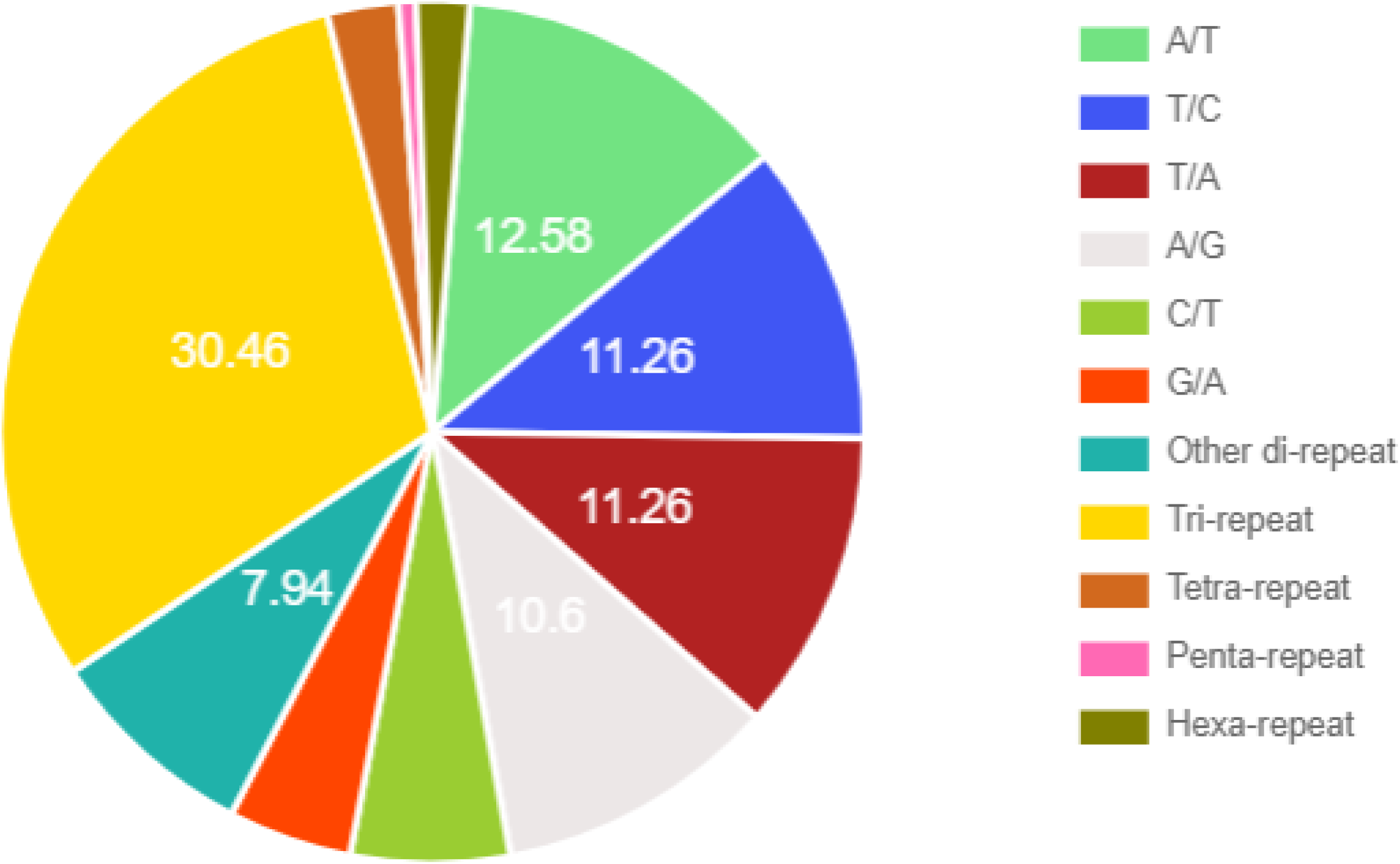
Distribution of the different repeat motifs that were detected in the mango RGA transcriptome.

**Figure 8.**
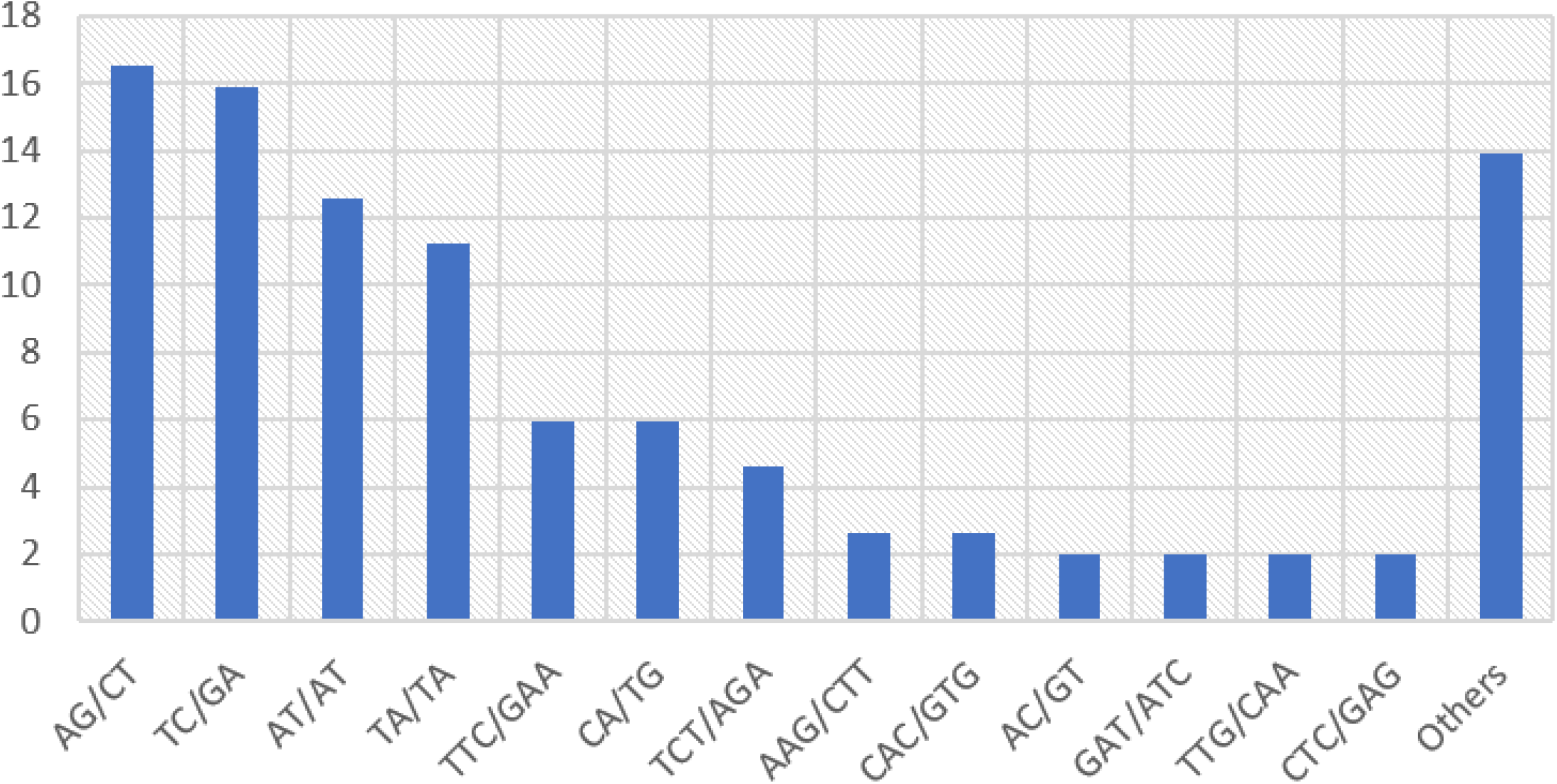
Paired-repeat SSR motif distribution of the mango RGA transcriptome.

**Figure 9.**
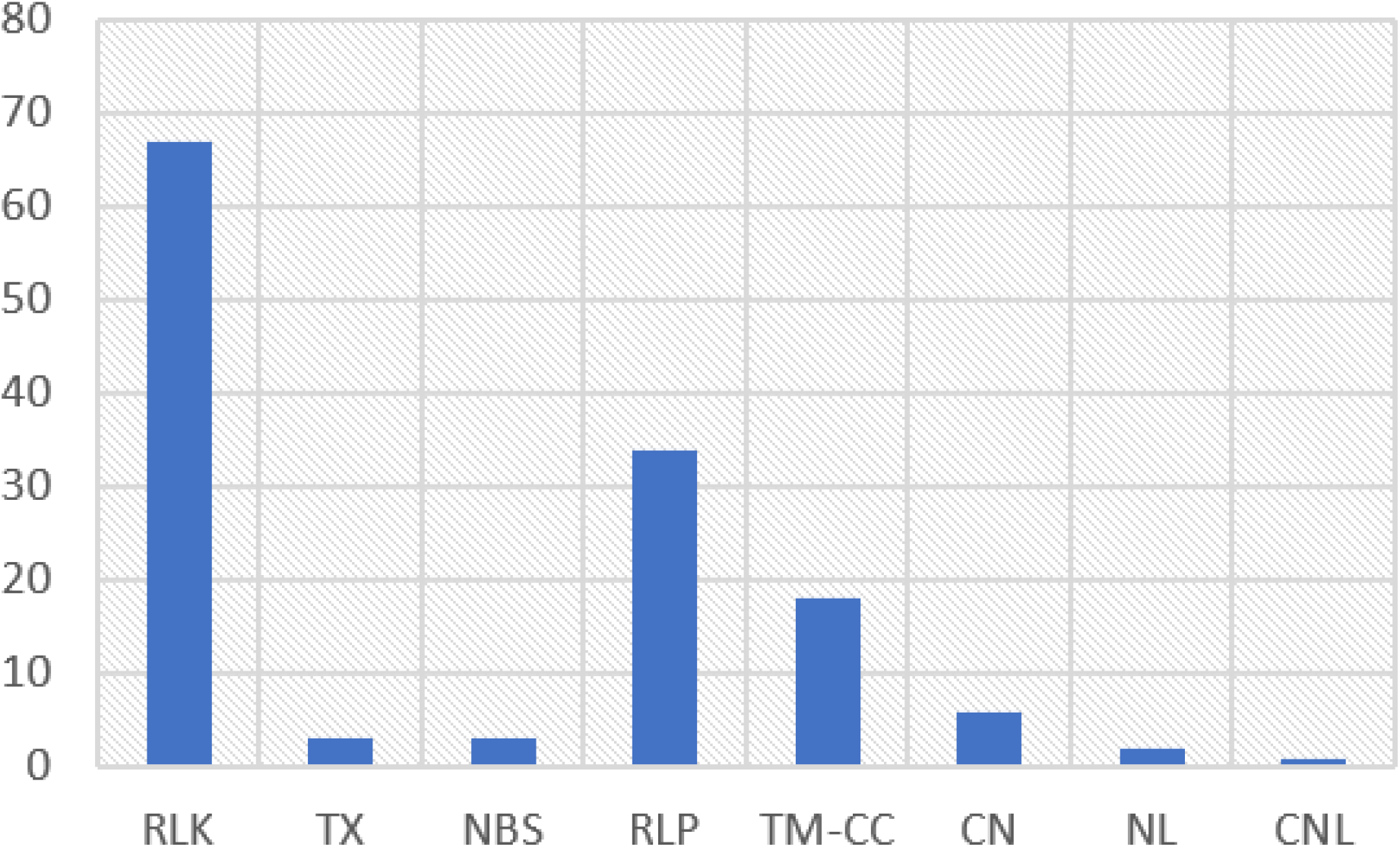
Distribution of EST-SSR markers designed across the different RGA domains identified in *de novo* mango transcriptomes.

Similar *in silico* approaches in EST-SSR marker development were performed by other studies focusing on other plants, such as bamboo (Cai et al., 2019), mint (Kumar et al., 2015), castor (Thatikunta et al., 2016), Ethiopian potato (Gadissa et al., 2018) and *Elymus* species (Zhang et al., 2019). The EST-derived SSR markers generated in this study can be used for functional analysis of *R* genes, once verified by wet lab experiments. Moreover, these markers may also be used in genotyping, genetic diversity, linkage mapping, gene-based association studies and marker-assisted selection in mango breeding. The sequences of the designed forward and reverse primers derived from mango RGAs are found in Supplemental Table 4.

## Summary and Conclusions

In this paper, 747 core mango RGAs were identified and broadly categorized into four major families based on their conserved structural features: NBS-encoding proteins (155 proteins), RLPs (158), RLKs (362) and TM-CC (72). In addition, the mango RGAs have been functionally well-annotated revealing the involved biological processes, molecular functions and cellular components. This provides an important insight to the overall functional response of mango against insect pests and diseases. Moreover, the evolutionary relationships, diversity, and expression profiles across different mango varieties were presented as an invaluable reference in the design of effective framework for the actual mango resistance breeding programs. A marker resource, comprising of 134 RGA-linked EST-SSR markers, was also developed to aid in the discovery of associated RGA/s to a specific disease or insect pest through population genetics analyses (e.g., QTL mapping, association mapping). The basic information from the mango RGAs and the linked EST-SSR markers established in this paper will facilitate in the development of outstanding mango varieties with resistance to various pests and diseases.

## Acknowledgements

The authors would like to thank the Department of Science and Technology-Philippine Council for Agriculture, Aquatic and Natural Resources Research and Development (DOST-PCAARRD) for funding the project entitled “Characterization of ‘Carabao’ and other Mango Varieties with Resistance to Fruit Fly and Anthracnose”. Also, the University of the Philippines Los Baños for the core-funded project, “Genome-wide analysis of resistance gene analogs (RGAs) and development of RGA-linked DNA markers in Philippine Important Crops”.

